# Global transmission of *Salmonella enterica* serovars associated with historical pork trade via Euramerica centralized sourcing

**DOI:** 10.1101/2022.09.30.510286

**Authors:** Yilei Wu, Lin Zhong, Shengkai Li, Heng Li

## Abstract

Past decades have witnessed swine as the common sources of *Salmonella enterica* infections. Nevertheless, how *Salmonella enterica* entered food supply chains and caused foodborne outbreaks remains to be excluded. By analysis of 718 publicly *Salmonella enterica* isolates, we demonstrated that Euramerica act as the centralized sourcing for the global spread of *Salmonella enterica* which origins at the pinnacle of pork production. Specifically, the lineage of *Salmonella* Choleraesuis lend geographic support for their dispersal from Euramerica centralized origins. We used the generalized linear model to quantify the relative contributions of potential explanatory variables to the global transmission of *Salmonella enterica*. The global pork trade may act as the key driver for the geographic dispersal of *Salmonella enterica* over the world. In addition, the application of antimicrobials over the past 80 years may also exert a large impact on the evolution of *Salmonella enterica* genomics. Taken together, our findings provided the global lineage evolution of *Salmonella enterica*. Further investigation and potential intervention are needed to identify the global spread of *Salmonella enterica* over the world.

## Introduction

*Salmonella enterica* entered food supply chains and caused foodborne outbreaks via contaminated food, water or food-processing facilities, leading to tremendous life and economic losses worldwide (1). There are >2000 genetically independent natural populations in *Salmonella enterica*, which were often recognized by their serovars or eBGs (ST complexes) based on the multi-locus sequence typing (MLST) scheme. Most *Salmonella enterica* populations are considered “generalists” because they were found in various sources and hosts, while some were “specialists” that lived in a limited range of sources or have been adapted to a single host (2). Except for some special cases, it is still largely unknown how specialists have been transmitted internationally.

Pork and pigs are common sources of *Salmonella enterica* infections, counting for ∼31.1% of Salmonellosis and 9.3% of disease outbreaks in the European Union (3). A wide spectrum of *Salmonella enterica* serovars has been identified in all procedures of pig production. These include generalists like serovar Typhimurium and its monophasic variants, and a set of specialists including Choleraesuis, which allowed investigation of the international transmissions of bacterial pathogens in the sector specifically (4).

First isolated in 1886, *Salmonella enterica* serovar Choleraesuis was the first recognized animal pathogenic salmonellae. It was prevalent in pigs globally before the 1995s, leading to tremendous economic losses in the sector, but experienced a continuous decline in population size till now (5). Besides, Choleraesuis also causes systematic infections in humans and was the 10th most isolated serovars from humans in Southeast Asia from 1993 to 2002 (6). Phylogenetically, Choleraesuis belonged to a group of ancient *Salmonella*, namely the Para C Lineage. The Para C Lineage probably became a specialist over 4000 years ago and evolved the host specificities to humans (Paratyphi C), swine (Typhisuis) or both (Choleraesuis) (7). Lots of gene disruptions and metabolic recrafts were evidenced during the process. However, it is unclear whether the host adaption has stopped, or is still an ongoing process nowadays.

The evolution of the Para C lineage has been closely interacting with the pig domestication process. European Pigs were originally migrated from Mideast during the Neolithic, and later re-domesticated from local wild boar (8). During the industrial revolution, European pigs interbred with Asian pigs to form modern breeds (9). With the inventions of antimicrobials and intensive pig farming, Europe became the center of a global trading system for not only pork but also pig breeds (9). Previous studies have shown that wild boars were important mediates for regional transmission of serovar Choleraesuis (10). However, it is still unclear how the serovar has been transmitted to other regions worldwide, and how modern agriculture recrafted the genotypes and phenotypes of this pathogen.

Through the analysis of 718 publicly available isolates, we attempted to comprehend and clarify the continental and transcontinental transmission pathways of *Salmonella enterica* throughout the previous 200 years. Geographical information with additional analysis of swine trade and antimicrobial resistance genes was additionally performed in the present study.

## Methods

### Genome data retrieval and processing

A total of 893 Salmonella isolates of the Para C lineage were analyzed in this study. 877 genomes were assembled by the Enterobase *Salmonella* database (11). The Chinese Center for Disease Control and Prevention provided an additional 16 novel *Salmonella* Choleraesuis isolates from mainland China. These genomes were isolated between -3242 and 2020 and originated from 51 countries. The specific serotyping, geographic information and source details were extracted from the metadata files. Genomic reads were trimmed and assembled using EToKi (https://github.com/zheminzhou/EToKi). After filtering by metadata, the collection of 893 Para C lineage sequences and two collections of 718 and 583 *Salmonella* Choleraesuis sequences were carried out for their subsequent analysis (583 genomes have no ancient isolates and only contain isolates with completed information).

### Temporal signal and randomization test

Due to the significance of temporal structure for evolutionary analysis, we identified weak temporal signals of *Salmonella* Choleraesuis isolates by a root-to-tip distance corresponding to their isolation dates using TempEst v1.5.3, TreeTime v0.8.5 and BactDating v1.1 (12–14). The date-randomization test was conducted by the TreeTime subcommand clock. We simulated multiple random-dated datasets compared with the original dataset by the substitution rate and coefficient of determination. The outcome showed a relative strong temporal structure, as the substitution rate of the real dataset did not fall in 95% of the credible interval of the substitution rates of replicated datasets.

### Phylogenetic analysis and Date estimation of the MRCA of *Salmonella* Choleraesuis

EToKi was used for multiple sequence alignments, removal of recombination regions (RecHMM and RecFilter) and construction of a phylogenetic tree for 718 Choleraesuis genomes based on SNP. For each clade, we assigned a synonymous mutation as a marker. Based on the SNP alignment of 583 genomes of Choleraesuis collection, IQ-Tree v1.6.12 built an SNP tree (15). Maximum-likelihood tree of *Salmonella* Choleraesuis was built by BEAST and BactDating (12,16), which both performed Bayesian inference of ancestral dates on the nodes of the SNP tree. Tree visualization was done using iTol (17). The tMRCA of 718 Choleraesuis collection was measured by BEAST (374 BC, 501-256 BC). The tMRCA for the collection of 583 genomes (without the ancient genomes) was initially calculated by TreeTime (757 CE), and the result was also within the 95% confidence interval of the time estimated by BactDating (722 CE, -761.96-1405.33 CE).

### Trade data and transmissions of multiple serovars related to swine

Data for the international trade of live poultry and pork were obtained from Harvard Dataverse (https://dataverse.harvard.edu). Historical trade values were adjusted for inflation and expressed in constant U.S. dollars. Scripts using Basemap Toolkit v1.2.0 in Python 3.5 visualized the intercontinental trade of pork and live poultry and transmission of the whole nine serovars related to swine. Transmission of each serovar was performed by chord diagrams using circlize packages in R.

### General linear model (GLM) extension of Bayesian phylogeographic inference

Potential explanatory variables for Salmonella Enterica dispersal were quantitatively assessed using the generalized linear model extension of Bayesian phylogeographic inference as previously described (18). Variables tested include the trade of pork, pig breeders and live pigs. BEAST analyses were performed using the Bayesian skyline model in combination with a relaxed log-normal molecular clock (16).

### Antibiotics and mobile elements detection

Resfinder v4.0 was used to detect antibiotic resistance genes and mobile elements (19). The patterns between antibiotics and mobile elements were detected by custom scripts.

## Results

### Phylodynamics of *Salmonella enterica* serovar Choleraesuis lends geographic support for their dispersal from Euramerica’s centralized origins

To further investigate the global transmission of *Salmonella enterica* serovar Choleraesuis, we performed the genomic data of 718 *Salmonella* Choleraesuis isolated from 40 countries between 1935 and 2020 that were available at EnteroBase (Figure 1A). The population structure and evolutionary factors were analyzed by identification of a phylogenetic tree with 19,294 chromosomal single-nucleotide polymorphisms (SNPs) after the removal of recombination. Results showed that the majority of the strains were isolated from West European countries (293/718; 40.8%), followed by North American countries (124/718; 17.3%). The global *Salmonella* Choleraesuis was distributed into three major clades 1, 2, and 3 with 19 subclades being identified according to the synonymous mutation.

**Figure 1.**
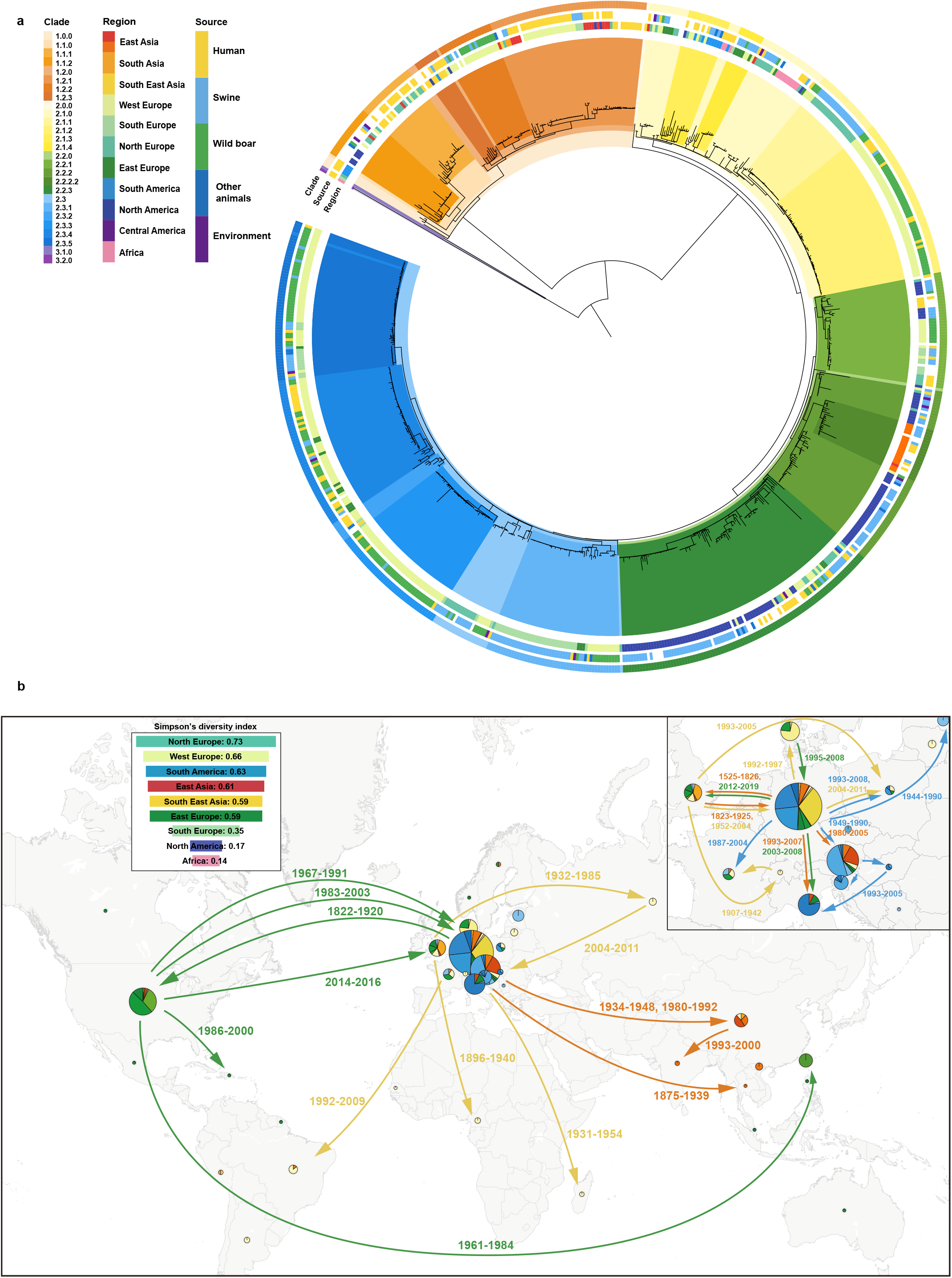
The global transmission of *Salmonella enterica* serovars associated with pork trade via Euramerica centralized sourcing.

Clade 1 included Choleraesuis strains from Southeast Asia and West Europe (68/134; 50.7%). Clade 2 contained the majority of the global genomes (582/718; 81.1%) with three subclades 2.1, 2.2, and 2.3. Subclade 2.1 included isolates from swine and wild boar in Northern and Western Europe, as well as human samples from Africa, Southern America, and other parts of Europe. Subclade 2.2 consisted of the isolates transmitted between Northern America, Europe, and East Asia. Approximately three-fifths of the genomes in Clade 2.2 were isolated from swine, while only nearly one-fifth from humans. Notably, the Taiwan subclade contained 18 genomes from 1998 to 2015 belonging to the US clade, suggesting the potential cross-border transmission. Additionally, Clade 2.3 were Choleraesuis mainly isolated within Europe and Clade 3 only consisted of 2 genomes isolated from Italy and Algeria (Figure 1A).

The present phylodynamics of *Salmonella* Choleraesuis lineages lend geographic support for their dispersal from Euramerica’s centralized origins. The evolution and divergence period of the worldwide *Salmonella* Choleraesuis across several internal clades were examined using a combination of Bayesian inference with ancestral and country information. Results showed that the coalescent date of Choleraesuis could be traced back 1,300 years ago, with Western Europe and North America being the top two global centers for porcine Choleraesuis isolates that subclade 2.1 had spread successively to Africa and Southern America with Europe as the center (Figure 1A). Subclade 2.2 was centered in the US and had mainly spread to Western and Northern Europe in the last 50 years.

### International porcine trade is concordant with the global transmission of additional *Salmonella enterica* serotypes

Historical data on international porcine trade are publicly available at Harvard Dataverse. We extracted the pork trade data from 1990 to 2021 and find that the past three decades had witnessed a domination trade with European and North American countries as exporter regions (Figure 1B). East Asian countries such as China and Japan import large quantities of pork from Europe (Green) and the United States (Blue). Similarly, in comparison between porcine trade data and Salmonella spread, we found that international porcine trade was concordant with the global transmission of additional *Salmonella enterica* serotypes via Euramerica centralized sourcing (Figure 1B). Then we focus on the *Salmonella enterica* serovar Choleraesuis. Based on the analysis of average transmission frequency, we found that swine was a significant issue in bacterial transmission. Swine also played a role in the transmission of *Salmonella* Choleraesuis between food, environment, humans and other livestock (Figure 1B).

We used the generalized linear model (GLM) extension of Bayesian phylogeographic inference to quantify the relative contributions of potential explanatory variables to the global transmission of *Salmonella enterica*. The international trade of raw pork may present as the driver for the spread of the Global lineage to the sampled countries as suggested by the GLM analysis (R^2^=0.5835). However, other factors such as pig breeders (R^2^=0.1491) and live pigs (R^2^=0.0713) were not significant to the bacterial transmission. Taken together, our quantitative assessment identified the global pork trade as the key driver for the geographic dispersal of *Salmonella enterica* over the world.

### Antimicrobial consumption in swine may serve as one of the key points in the evolution of *Salmonella enterica*

The agricultural use of antimicrobials after intensive pig farming may play the role in the evolution of antimicrobial-resistant *Salmonella enterica* strains. The antimicrobial resistance in *Salmonella* Choleraesuis showed an increasing and decreasing trend in the past hundred years. Lineage 1 witnessed an increase in antimicrobial resistance in the 1940s and the following decade. Antimicrobial resistance in Choleraesuis lineage 2 showed an increasing trend in the early 1960s. Following that, the mean value peaked around 2000 and declined rapidly over the next 20 years with the injection of the Avirulent vaccine.

Specific antimicrobial resistance genes were screened among 718 *Salmonella* Choleraesuis isolates. Resistance to sulfonamides, aminoglycoside, tetracycline, and ampicillin was higher detected in *Salmonella* Choleraesuis strains. The East Asian isolates showed 36% (54/151) of the global distribution of resistance to quinolones, whereas other countries such as the UK, US, and China shared relatively high levels of antimicrobial resistance, suggesting that the applicartion of antimicrobials over the past 80 years exert a large impact on the evolution of *Salmonella enterica* genomics.

## Discussion

The traditional theories pronounced that geographical location played a dominant role in the occurrence and evolution of bacteria (20). However, the case between China and Taiwan showed that the accelerating global trade in livestock may also be an important factor in the spread of bacteria. Traceback investigations combining trade data and pathogen genomes have resolved recent transmissions of *Salmonella enterica* at the scale of individual outbreaks in Europe (21,22). Here combined with the whole-genome sequencing, we elucidated the genomic epidemiology study of *Salmonella* Choleraesuis interwoven with the intercontinental trade during the past decades.

We thoroughly reviewed the transmission of *Salmonella* Choleraesuis during the historical timeline. The pre-industrial culture dating back to the year 1500, the industrial revolution from 1750 to 1940, and the post-antibiotic era were the three stages of the transmission model that were shown. Clade 1 was first predicted to emerge and spread over Europe before this lineage was then transmitted to Asia and other regions after the industrial revolution around 1850 to 1900. Simultaneously, Clade 2 began to spread in the industrial era with Europe and America as centralized origins. Isolates from North America were located in clusters with those from East Asia and Europe. Subclade 2.2 mainly consisted of trade between the US and Europe, there were still some isolates from Taiwan that reflected the different swine sources between mainland and Taiwan in the 1970s. The phylogeographic analysis showed that the transmission of *Salmonella* Choleraesuis between countries was concordant with the trading network of pigs.

The inclusion of several *Salmonella enterica* serovars has led to a historical conundrum about the pandemic’s origins (23). We reported ancestry status reconstructions suggested a common European and American ancestry for *Salmonella* Choleraesuis transmission across global lineages. Highly diversified *Salmonella* Choleraesuis populations in swine would come from the unrelated and dispersed origins of the serotype on several continents, representing the serotype’s complex worldwide evolutionary history. Second, phylodynamic analyses provide geographic support to the hypothesized central origins. The global lineage appears to be a direct consequence of swine-mediated *Salmonella* transmission, its population number increased considerably during the industrial revolution and the post-antibiotic era. According to estimates, the emergence of the European Subclade 2.3 occurred earlier than the start of pedigree breeding. This clade probably corresponds to early swine strains circulating in Europe at the time of the emergence of industrialized swine farming (24).

The *Salmonella* Choleraesuis pathogens are mainly divided into two clusters with four clades in the present study. Except intra-European Subclade 2.3, the other clades reflect the pathogen transmission from North Europe and the US to other regions in Europe, Asia, and Africa. According to the Simpson diversity index, commerce between geographically adjacent regions like Northern and Western Europe was more common, which will enhance the diversity of Choleraesuis pathogens and aid in their adaptability. From 1950 to 2000, mainland China imported breeds of pigs such as Yorkshires and Landraces from Europe, whilst Taiwan bought goods from the US between 1960 and 1990 (25,26). Importantly, trace back investigation of *Salmonella* isolates, with 16 new pathogen genomes from mainland China, revealed the intercontinental transmission between East Asia and Western countries.

Another confounding factor to our findings is antimicrobial overuse which should be prevented and limited by humans due to the multidrug resistance displayed by isolates from the Para C lineage. In the clinic, infections are usually treated with sulfamethoxazole/trimethoprim, ciprofloxacin, or cephalosporins, which were associated with the antimicrobial resistance that has been increasing in Salmonella since the 1980s (27). One factor influencing the spread of antimicrobial resistance is horizontal gene transfer (HGT). Mobile genetic elements, including plasmids, and transposons, have featured prominently as agents of HGT associated with AR in Enterobacteriaceae, including *Salmonella* and facilitate HGT via conjugation, transduction, and transformation, respectively (28). Notably, the colistin-resistant gene *mcr*-3, commonly detected in swine, was found in six Choleraesuis strains isolated from the human infection in China and the UK via global screening, suggesting the urgent need for surveillance of this resistance (29,30).

In conclusion, our findings shed light on the global lineage evolution of *Salmonella enterica* in terms of intercontinental transmission, exogenous and endogenous factors such as the emergence and accumulation of antimicrobial resistance. Despite decades of significant progress on *Salmonella* control in swine, the evidence provided in our study warrants further investigation and potential intervention into the global spread of *Salmonella enterica* from centralized origins at the pinnacle of pork production.

## Ethics approval and consent to participate

Not applicable

## Consent for publication

The Author hereby consents to publication of the Work in any and all publications.

## Availability of data and materials

The datasets used and/or analysed during the current study available from the corresponding author on reasonable request.

## Authors’ contributions

YW, HL wrote the main manuscript text and LZ, SL prepared the data. YW, HL analysed the statistical data. All authors reviewed the manuscript.

## Acknowledgements

Not applicable

